# Single cell transcriptomic analysis revealed long-lasting adverse effects of prenatal tamoxifen administration on neurogenesis in prenatal and adult brains

**DOI:** 10.1101/811893

**Authors:** Chia-Ming Lee, Liqiang Zhou, Jiping Liu, Jiayu Shi, Yanan Geng, Jiaruo Wang, Xinjie Su, Nicholas Barad, Junbang Wang, Yi E. Sun, Quan Lin

**Affiliations:** Department of Psychiatry and Behavioral Sciences, Intellectual Development and Disabilities Research Center, University of California, Los Angeles, CA 90095; Stem Cell Translational Research Center, Tongji Hospital, Tongji University School of Medicine, Shanghai, China 200092

## Abstract

CreER/LoxP system has enabled precise gene manipulation in distinct cell subpopulations at any specific time point upon tamoxifen (TAM) administration. This system is widely accepted to track neural lineages and study gene functions. We have observed prenatal TAM treatment caused high rate of delayed delivery and mortality of pups. These substances could promote undesired results, leading to data misinterpretation. Here, we report that TAM administration during early stages of cortical neurogenesis promoted precocious neural differentiation, while inhibited neural progenitor cell (NPC) proliferation. The TAM-induced inhibition of NPC proliferation led to deficits in cortical neurogenesis, dendritic morphogenesis, and cortical patterning in neonatal and postnatal offspring. Mechanistically, single cell RNA sequencing (scRNA-seq) analysis combined with *in vivo* and *in vitro* assays showed TAM could exert these drastic effects mainly through dysregulating the expression of *Dmrta2* and *Wnt8b.* In adult mice, administration of TAM significantly attenuated NPC proliferation in both the subventricular zone and the dentate gyrus. This study revealed the cellular and molecular mechanisms for the adverse effects of prenatal tamoxifen administration on corticogenesis, suggesting that tamoxifen-induced CreER/LoxP system may not be suitable for neural lineage tracing and genetic manipulation studies in both embryonic and adult brains.

**Significant:** For the first time, our study revealed the molecular mechanisms underlying tamoxifen activities on cortical development. This study also clearly showed that care must be taken when using tamoxifen-induced CreER/LoxP system for neural lineage tracing and genetic manipulation studies.

## Introduction

Tamoxifen (TAM)-inducible CreER/LoxP system is one of the most widely used genetic tools that have enabled precise gene manipulation in distinct cell subpopulations at any specific time point which is known as temporally and spatially controllable gene expression. Besides gene knockout, CreER/LoxP technology has enabled labeling of any cell types for genetic fate-mapping study (1-4).

TAM, a selective estrogen receptor modulator (SERM), is considered a pioneering and commonly used drug to treat estrogen receptor positive breast cancer (5). In humans, a study based on AstraZeneca Safety Database suggested that TAM therapy of breast cancer during pregnancy is tightly associated with spontaneous abortions, fetal defects, and congenital malformations (6). To avoid significant birth defects, TAM administration needs to be discontinued during pregnancy in breast cancer patients (7-9). In rodents, adverse effects of TAM on prenatal and postnatal mice have been reported by a considerable number of studies. Empirically, administration of TAM or its active metabolite, 4-hydroxytamoxifen (4-OH-TAM) by either gavage (~1.5-10mg TAM/mouse) or intraperitoneal (IP) injections (~750μg-1.5mg TAM/mouse) to pregnant mice led to embryonic lethality and/or dystocia. To obtain postnatal offspring, it is frequently required that a caesarian section is performed followed by fostering (10-15).

Our initial study showed that administration of TAM to pregnant mice at E10 greatly reduced the size of the cerebral hemispheres and the olfactory bulbs, enlarged the lateral ventricle, and thinned the cortical plate (Fig. 1A), suggesting prenatal TAM exposure has a drastic impact on cortical neurogenesis. We delivered TAM to more than 70 litters of three different mouse lines by IP injection (i.e. C57BL/6, 129S6/C57BL/6, & CD1). TAM dosages tested per pregnant dam were 500µg (5 litters), 750µg (≥50 litters), 1mg (5 litters), and 2mg (≥10 litters). The delayed delivery and/or mortality of pups were always observed in litters with TAM dosage of 750µg/animal and above.

**Figure 1.**
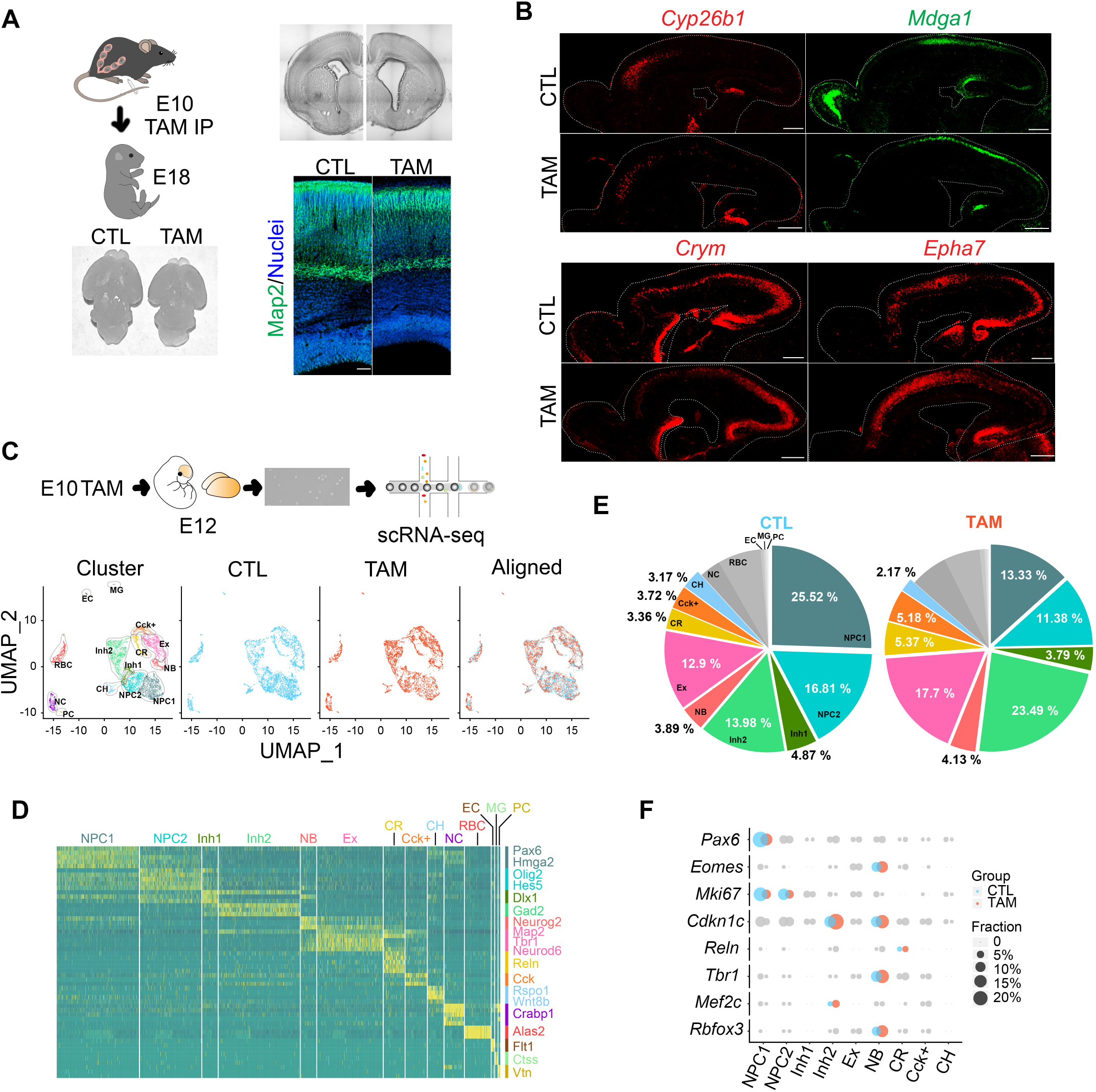
Prenatal TAM exposure impaired cortical neurogenesis. **(A)** TAM was intraperitoneally injected into pregnant mice at E10 (750µg/animal). The brain was removed at E18. TAM treatment reduced the size of the cerebral hemispheres and the olfactory bulbs. Nissl staining showed an enlargement of the lateral ventricles (upper right panel). Map2 immunostaining shows TAM treatment at E10 dramatically thinned cortical plate at E18 (lower right panel). CTL, oil control; TAM, tamoxifen treated; Map2, microtubule associated protein 2. Scale bar, 50μm. **(B)** Prenatal TAM exposure at E10 altered cortical area patterning at E18 in the motor cortex (*Cyp26b1*, *Epha7*), the somatosensory cortex (*Mdga1*), and the visual cortex (*Crym*, *Epha7*). Scale bar, 400μm. *Cyp26b1*, cytochrome P450 family 26 subfamily b member 1; *Epha7*, EPH receptor a7; *Mdga1*, MAM domain containing glycosylphosphatidylinositol anchor 1; *Crym*, Crystallin Mu. **(C)** Schematic overview of single cell isolation and 10X Genomics scRNA-seq. TAM, tamoxifen IP injection at E10. Visualization of single cell data by uniform manifold approximation and projection (UMAP) plot of clustering of E12 cells. NPC1, pallial neural progenitor cells; NPC2, subpallial neural progenitor cells; Inh1, inhibitory neurons (mainly derived from the medial ganglionic eminence (MGE)); Inh2, inhibitory neurons (mainly derived from the lateral ganglionic eminence (LGE) or the caudal ganglionic eminence (CGE)); Ex, excitatory neurons; NB, neuroblasts; CR, Cajal-Retzius cells; CH, cortical hem cells; NC, neural crest cells; RBC, red blood cells; EC, endothelial cells; MG, microglia; PC, pericytes. **(D)** Heatmap shows top cell-type-specific genes expressed in clusters defined in (C). **(E)** Pie chart shows major cell-type percentages. **(F)** Patterning analysis of cell fate and cell cycle markers. Circles represent fraction of positive cells in each cluster. Pax6, paired box 6; Eomes, eomesodermin; Mki67, marker of proliferation Ki-67; Cdkn1c, cyclin dependent kinase inhibitor 1C; Reln, reelin; Tbr1, T-box, brain 1; Mef2c, myocyte enhancer factor 2C; Rbfox3, RNA binding Fox-1 homolog 3.

To reveal the adverse effects of TAM on cortical neurogenesis, we carried out scRNA-seq analysis. Combined with in situ hybridization (ISH) and immunostaining assays, we report the biphasic effects of TAM treatment during cortical neurogenesis. On the one hand, TAM promoted precocious cortical neurogenesis. On the other hand, it blocked NPC proliferation and thus ultimately resulted in impairment of cortical neurogenesis in postnatal offspring. Mechanistically, TAM could exert these drastic effects through downregulating the expression of *Dmrta2* and its effector *Hes1*, which have been shown to play critical roles in NPC fate maintenance and neural differentiation (16, 17). Moreover, in TAM treated brains, misexpression of *Wnt8b* in the cortical hem and downregulation of Wnt and Bmp receptors, *Lrp6* and *Acvr2b*, respectively in the cortex may cause cortical patterning defects. In addition, we showed that TAM administration at E8.5 severely impaired neural dendritic morphogenesis and blocked gliogenesis in postnatal offspring. In adult mice, TAM treatment dramatically attenuated neural stem cell proliferation in the subventricular zone (SVZ) and the dentate gyrus (DG), the two areas where adult neurogenesis occurs.

## Results

### Prenatal TAM administration impaired neurogenesis and cortical organization in perinatal offspring

It has been shown that the highest dose tolerance for a single prenatal (E7.5-10.5) TAM IP injection without significant lethality at term is about 1mg (~30-60μg TAM per gram body weight) (Hayashi and McMahon, 2002; Paul S. Danielian, 1998). Our preliminary study showed administration of 500µg TAM to prominin 1 promoter drove CreER reporter mice (Prom1CreER/floxed ZsGreen) at E8.5 and E10 did not elicit detectable green fluorescent protein signal at E18. Therefore, to study the effects of TAM on prenatal cortical development, we chose to administrate 750µg of TAM per pregnant dam (~25-30μg TAM per gram body weight).

Administration of TAM to pregnant mice at E10 reduced the size of the cerebral hemispheres at E18, the time when cortical neurogenesis is largely completed (18) (Fig. 1A). We found administration of TAM at E10 dysregulated the expression of multiple cortical patterning genes in the motor cortex (*Cyp26b1*, *Epha7*), the somatosensory cortex (*Mdga1*), and the visual cortex (*Crym*, *Epha7*) in E18 offspring (Fig. 1B). TAM treatment somewhat reduced visual cortex marker gene expression, and also expanded the somatosensory gene *Mdga1* expression posteriorly, implying cortical patterning deficits in TAM treated offspring. To reveal the effects of TAM on corticogenesis thoroughly, we carried out scRNA-seq analysis.

TAM was delivered to the pregnant dam at early stage of cortical neurogenesis (E10) by IP injected. At E12, the left and the right hemispheres were pooled and subjected to droplet-based scRNA-seq analysis (10X Genomics) (Fig. 1C). We applied quality filter and integrated analysis to merge control (CTL) and TAM datasets with total of 8,276 cells and 17,071 genes using the Seurat package (19, 20). Based on uniform manifold approximation and projection (UMAP) analysis, we segregated the cells into 14 clusters (Fig. 1C). According to genes expression enrichment, we annotated the 14 clusters using major cell type markers of each cluster. There are neural progenitor cells (NPC1, *Pax6* and NPC2, *Olig2*), inhibitory neurons (Inh1, *Dlx1* and Inh2, *Gad2*), neuroblasts (NB, *Neurog2*), excitatory neurons (Ex, *Neurod6*), Cajal-Retzius cells (CR, *Reln*), Cck+ neurons (*Cck*), cortical hem neurons (CH, *Rspo1*), neural crest cells (NC, *Crabp1*), red blood cells (RBC, *Alas2*), endothelial cells (EC, *Flt1*), microglia (MG, *Ctss*), and pericytes (PC, *Vtn*) (Fig. 1C, D). The scRNA-seq data showed that prenatal TAM exposure greatly reduced total numbers of pallial and subpallial NPCs (NPC1&2, smalt blue &. blue color) by 15.62%, while caused a total of 16.7% increase in both cortical (Ex, pink color) and subcortical neurons (Inh1&2, green color), CR (dark yellow color), and Cck+ neurons (orange color) (Fig. 1E). Prenatal TAM treatment also reduced the number of cells in one of the major brain patterning centers, the cortical hem (light blue color) (21). We further analyzed ratios of progenitor and postmitotic cell clusters using cell fate markers, Pax6, Eomes (Tbr2), Reln, Tbr1, Mef2c, and Rbfox3 (NeuN), as well as cell cycle related genes, Mki67 and cyclin dependent kinase inhibitor 1C (Cdkn1c). TAM notably reduced the number of progenitors in the pallial VZ (Pax6+) but increased the number of SVZ cells (Eomes+) and postmitotic neurons (Rbfox3+), including superficial layer (Reln+, Mef2c+) and deep layer (Tbr1+) neurons. In TAM treated brains, the number of proliferating cells (Mki67+) was dramatically reduced, and there were more cells expressing the cell cycle G1-phase marker, Cdkn1c (Fig. 1F).

In summary, our scRNA-seq analysis suggested that prenatal TAM exposure inhibits VZ progenitor proliferation, while increasing the number of SVZ progenitor cells and promoting precocious neuronal differentiation. To investigate the biphasic effects of TAM, we sought to study the influences of prenatal TAM administration on cortical neurogenesis at different neurogenic stages and analyze its long-lasting effects in postnatal offspring.

### Prenatal TAM exposure inhibited VZ progenitor proliferation

To validate the initial scRNA-seq analysis, we first measured the total number of NPCs in the cortex using the VZ and the SVZ markers, Pax6 and Emoes, respectively. The ISH and immunostaining assays showed that in E10 TAM treated brains (750μg TAM/pregnant dam), the number of Pax6+ NPCs in the VZ was significantly reduced while Eomes+ NPCs in the SVZ was slightly increased (Fig. 2A, B, Sup. Fig. 1A). Second, bromodeoxyuridine (BrdU) labeling combined with Mki67 immunostaining showed that TAM exposure at E10 and E13 greatly reduced the number of cells in the cell cycle (Mki67+) and in the S-phase (BrdU+) (Fig. 2C, Sup. Fig. 1B). Third, we measured cell cycle S-phase length using a 5-ethynyl-2′-deoxyuridine (EdU)/BrdU double labeling paradigm (22) (Fig. 2D). Cells that left the S-phase during the labeling period (2hr) were labeled with EdU only (green color). Cells in S-phase were BrdU labeled (red color). The EdU/BrdU double labeling assay showed a prolonged S-phase in TAM-treated, proliferating neural progenitors (Fig. 2D). Fourth, we assessed the number of dividing cells (M-phase) in the cortex using phospho-Histone H3 (pH3) antibody (23). We found there were more basal pH3+ cells in TAM treated brains than the controls, which may cause transient increase of Eomes+ cells. Contrary to its effect on basal dividing cells, TAM treatment reduced the number of pH3-labeled, apical dividing cells in the cortex (Fig. 2E), which may cause reduction of Pax6+ NPCs and lead to neurogenesis impairment later in life (Fig.1A). Finally, we found TAM treatment at early and mid-stage of cortical development quickly increased the number of Cdkn1c positive cells (Fig. 2F, Sup. Fig. 1C). The cell cycle arrest molecule, Cdkn1c (P57Kip2) is a strong inhibitor of several G1 cyclin/Cdk (Cyclin-dependent kinases) complexes and a negative regulator of cell proliferation (24, 25). Increased number of cells in the G1-phase suggests an early cell cycle withdraw and subsequent advanced cortical neuronal differentiation. Taken together, it appears that in cortical NPCs, TAM administration disrupts cell cycle progression which in turn inhibits NPC proliferation. In the meantime, TAM promotes transient SVZ progenitor proliferation followed by early cell cycle withdraw, which leads to precocious neural differentiation.

**Figure 2.**
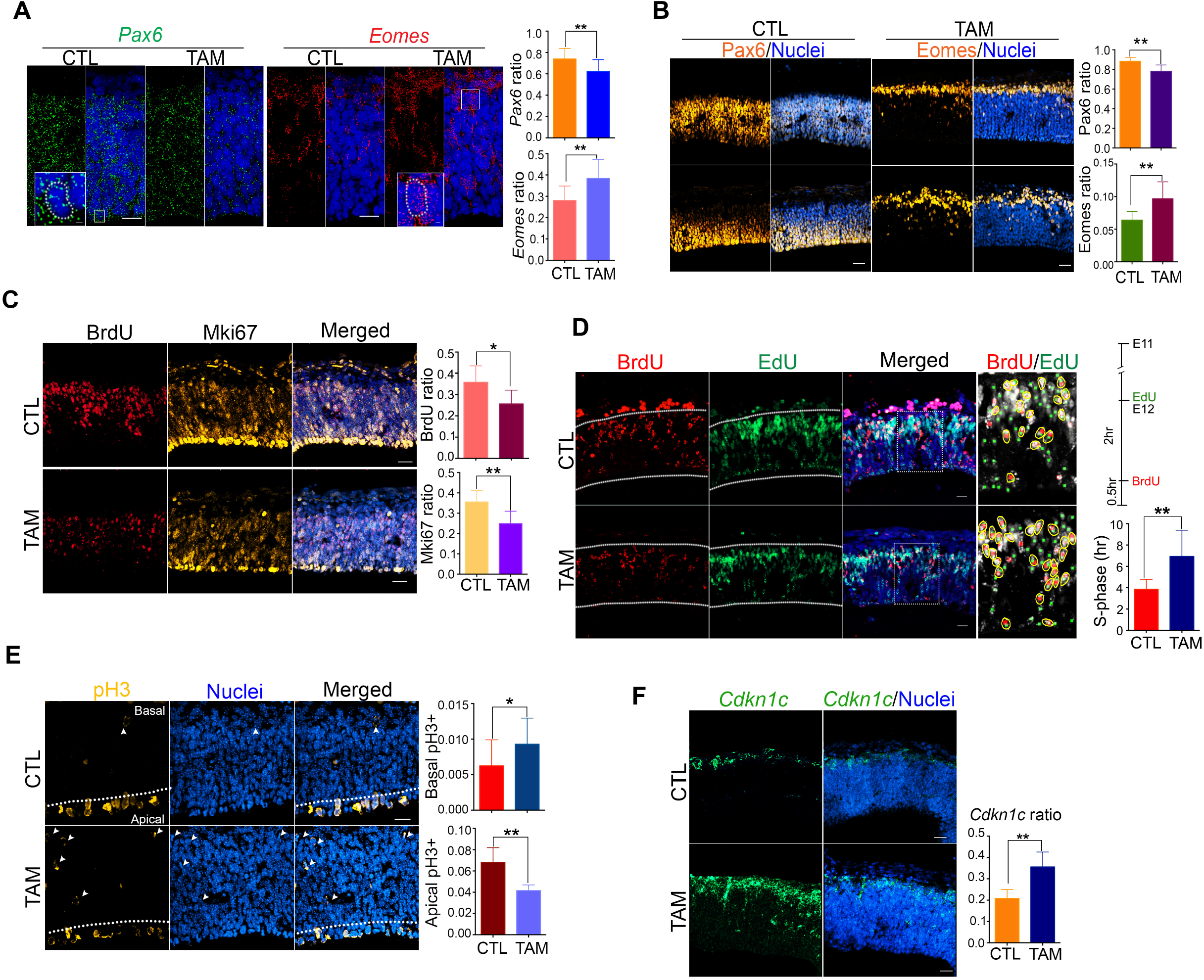
Prenatal TAM administration inhibited neural progenitor proliferation. **(A)** In situ hybridization (ISH) shows administration of TAM at early brain development (E10) reduced the number of ventricular zone (VZ) progenitors (*Pax6*+), while increased the number of SVZ progenitors (*Eomes*+) (*Pax6*+ cells: p=0.0250; *Eomes*+ cells: p=0.0228, Mann Whitney test, CTL n=7, TAM n=14). Scale bar, 20μm. **(B)** In agreement with the ISH assay in (A), Pax6 and Eomes immunostaining shows TAM significantly inhibited VZ progenitor proliferation, while increasing the number of cells in the SVZ (Pax6+ cells: p<0.0001; Eomes+ cells: p=0.0005, Mann Whitney test, CTL n=14, TAM n=14). Scale bar, 30μm. **(C)** Administration of TAM at E10 dramatically reduced numbers of cells in S-phase (BrdU+) and total cells in the cell cycle (*Mki67*+) (BrdU, p=0.0264, Mki67, p=0.0007, Mann Whitney test, CTL n=19, TAM n=18). BrdU was injected to the pregnant dam at E12. Samples were collected 30min after BrdU administration. Scale bar, 30μm. **(D)** Kinetic analysis of S-phase cell cycle progression. Prenatal TAM exposure increased the S-phase length (P<0.0001, CTL n=20, TAM n=21). TAM was administrated at E11. EdU was injected at E12. BrdU was injected 2hr after EdU administration. The brains were collected 30min after EdU injection. The enlarged images on the right showed quantification of BrdU (red color) and EdU (green color) single or double positive cells. The Edu/BrdU double positive cells were labeled with yellow ovals. Scale bar, 20μm. **(E)** TAM administration increased the number of basal dividing cells (phospho-Histone H3+), which might increase number of Eomes+ SVZ cells and thus led to precocious neurogenesis (p=0.0236, Mann Whitney test, CTL n=5, TAM n=8). Scale bar, 20μm. **(F)** *Cdkn1c* ISH shows prenatal administration of TAM at E10 detained cells in G1-phase (p=0.0044, Mann Whitney test, CTL n=12, TAM n=12). Scale bar, 30μm. Error bars, standard deviation.

To rule out the possibility that reduced number of NPCs is due to TAM-induced apoptosis, we carried out terminal deoxynucleotidyl transferase dUTP Nick-End Labeling (TUNEL) assay. We did not detect increased apoptosis in E12 brains when TAM (2mg/animal) was administrated at E10 (Sup. Fig. 1D).

### Prenatal TAM exposure promoted precocious and transient neuronal differentiation

The scRNA-seq analysis showed an increased number of cells expressing postmitotic neuronal markers, such as *Reln*, *Rbfox3*, *Tbr1*, and *Mef2c* at E12 when TAM was administrated at E10 (Fig. 1F). There was an increased number of cells expressing cell cycle G1-phase gene *Cdkn1c* in TAM treated brains, which further suggested a precocious neural differentiation. In agreement with above evidence, the immunostaining assays showed that TAM exposure greatly promoted the production of not only early-born, Reln+ Cajal-Retzius cells and deep layer, Tbr1+ neurons (Fig. 3A, B), but also superficial layer Mef2c+ cells (Fig. 3C) (26). These results were further confirmed by ISH assay using a pan-neuronal marker, *Rbfox3* (Fig. 3D). Moreover, we also showed that TAM treatment at the peak of cortical neurogenesis (E13-14) dramatically increased Tbr1+ and Mef2c+ cells in the cortical plate at E14, while having little effect on the total number of Reln+ cells in the cortical marginal zone (MZ) (Sup. Fig. 2A). This may be because Reln+ cells are one type of the earliest born cortical neurons which differentiated before TAM administration (27).

**Figure 3.**
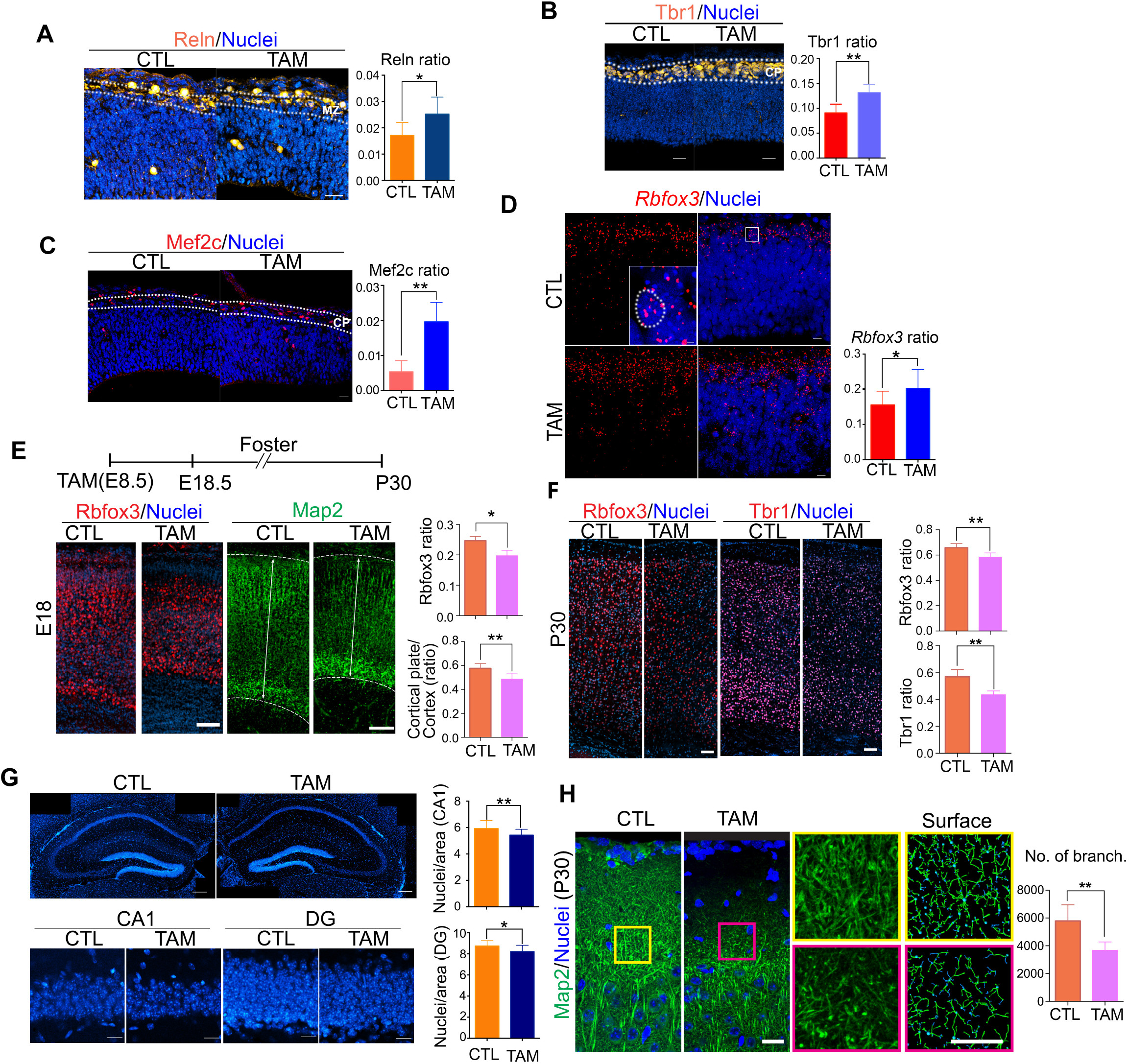
Prenatal TAM administration promotes precocious neurogenesis and has long-lasting effects on corticogenesis in postnatal offspring. **(A-D)** Administration of TAM at E10 increased numbers of Reelin (Reln) positive, Cajal–Retzius cells at E12 (p=0.0115, Mann-Whitney test, CTL n=7, Tam n=9, scale bar, 30μm), Tbr1+ early born neurons (p=0.0037, Mann-Whitney test, CTL n=8, TAM n=7, scale bar, 30μm), and superficial layer neurons (Mef2c+ cells) (p<0.0001, Mann-Whitney test, CTL n=20, TAM n=20; scale bar, 20μm). ISH assay shows prenatal TAM exposure at E10 transiently increased the total number of *Rbfox3*+ postmitotic neurons at E12 (p=0.0354, Mann-Whitney test, CTL n=10, TAM n=10). Scale bars, 10μm and 2μm (insert). **(E)** TAM was administrated at E8.5. Cortical neurons were significantly reduced in E18 brains (Rbfox3, p=0.0104, Mann-Whitney test, CTL n=15; TAM n=18; Map2, p<0.0001, Mann-Whitney test, CTL n=20; TAM n=35). Scale bar, 80μm. **(F)** Prenatal TAM exposure at E8.5 impaired cortical neurogenesis of P30 offspring (Rbfox3, p<0.0001, Mann-Whitney test, CTL n=15; TAM n=18; Tbr1, p=0.0044, Mann-Whitney test, CTL n=8; TAM n=8). Scale bar, 100μm. **(G)** Prenatal TAM administration at E8.5 perturbed neurogenesis in the hippocampus of P30 offspring. The CA1, p=0.0064, Mann-Whitney test, CTL n=18; TAM n=18. Scale bars, 200μm and 20μm. The DG, p=0.0141, Mann-Whitney test, CTL n=18; TAM n=18. Scale bars, 200μm and 20μm (zoomed in images). **(H)** TAM treatment decreased dendrite complexity in P30 offspring. Dendritic morphology was quantified using Imaris filament function, statistics denoted the number of branching points (p=0.0002, Mann-Whitney test, CTL n=9; TAM n=9). The right panels show enlarged the box areas in the left two panels and the surface model for dendrite quantification. Scale bar, 20μm. Error bars, standard deviation.

### The long-lasting effects of prenatal TAM exposure in postnatal offspring

To evaluate the potential long-lasting effects of TAM exposure on cortical neurogenesis, we injected TAM to pregnant dams preceding neural tube closure (E8.5) and examined cortical neurogenesis at E18 and at postnatal day 30 (P30). TAM treatment dramatically reduced the thickness of the cortical plate (Map2+) and the number of cortical neurons (Rbfox3+, Tbr1+) in both E18 and P30 offspring (Fig. 3E, F). In the hippocampus, prenatal TAM treatment also caused subtle but significant reduction of cell numbers in the CA1 and the DG (Fig. 3G). Moreover, administration of TAM at E8.5 severely impaired apical dendritic complexity in P30 brains, remarkably in the cortical marginal zone (Fig. 3H). Administration of TAM at E13 (the peak period of cortical neurogenesis) to an inbred strain (C57BL/6), an outbred mouse strain (Crl:CD1(ICR)), and a transgenic mouse line (Prom1CreER/ZsGreen (129S6/SvEvTac x C57BL/6NCrl)) also showed reduced number of cortical neurons (Sup. Fig. 2B, C), suggesting TAM treatment has long-lasting effects on cortical neurogenesis regardless of the genetic backgrounds.

In addition to cortical neurogenesis, in E8.5 TAM treated, P30 offspring, our data showed a significant reduction in the number of Olig2+ oligodendrocytes in the corpus callosum. The Cnp and Mbp staining also showed a thinner and disorganized axon track in the corpus callosum (Sup. Fig. 3A), suggesting prenatal TAM treatment impaired gliogenesis and/or dysregulated axon formation in postnatal offspring. We observed statistically non-significant reduction in the number of Gfap+ astrocytes in the corpus callosum, while Aldoc+ astrocytes remained unchanged in the cortex (Sup. Fig. 3B, C).

### Prenatal TAM exposure perturbed Wnt and BMP related signaling that has been shown to strongly correlate with neural progenitor maintenance, differentiation, and cortical areal organization

We have shown that prenatal TAM administration has broad effects from NPC proliferation, cell differentiation to neural circuit formation. To reveal underlying molecular mechanisms, we carried out gene regulatory network analysis of NPCs and cortical hem cells using bigSCale2 toolkit to identify hub genes that TAM employed to manipulate corticogenesis (28). We chose to analyze NPC1/2 and CH clusters because the tight association between VZ NPC proliferation and cortical neurogenesis. The cortical hem, one of main brain patterning organizers, has been shown to regulate cortical patterning (21, 29).

A total of 3,035 cells (CTL, 1,908 cells; TAM, 1,127 cells) were analyzed to generate a gene expression correlation network. We discovered four major network modules in CTL and three modules in TAM treated cells (Fig. 4A, left panels). Compared with the CTL, the TAM network showed lower edges to node density (CTL, 6.89; TAM, 4.92) and modularity (CTL, 0.61, four modules; TAM, 0.29, three modules), suggesting the CTL network preserved a strong intra-modular connectivity and higher gene expression heterogeneity. Gene ontology (GO) enrichment analysis showed the four modules in the CTL, comprising genes mainly related to forebrain development (e.g. *Nfib*, *Dmrta2*, *Emx2*, *Wnt8b*, *Dmrt3*, *Axin2*, *Tcf4*, *Meis2*, *Lmo1*), cell cycle progression (e.g. *Top2a*, *Ccnb1*, *Cenpf*, *Ctcf*, *Nucks1*), ribonucleoprotein complex biogenesis (e.g. *Ncl*, *Npm1*), and ribosome assembly and translation regulation (e.g. *Btf3, Rpl12, Rps5*) (Fig. 4A, right panels). Compared to CTL, TAM treatment reduced the GO enrichments to three modules that comprised genes mainly related to projection assembly and Wnt signaling (e.g. *Lmx1a*, *Axin2*, *Sostdc1*, *Wnt8b*), cell cycle progression (e.g. *Top2a*, *Ccnb1*, *Cenpf*), and ribosome assembly and translation regulation (e.g. *Ncl*, *Npm1*, *Rpl12*) (Fig. 4A, right panels). Closer inspection revealed the pagerank centrality of differentially expressed hub genes (orange-red color) reduced with random pattern in the gene regulatory network of TAM-treated NPCs and CH cells (Fig. 4A, left panels, orange-red color). The most striking example is the forebrain development module. The regulatory networks of the hub genes in this module (e.g. *Dmrta2*, *Wnt8b*, *Emx2*, *Dmrt3*) were greatly altered in TAM treated cells. For example, directly connected genes (nodes) (neighborhood of graph vertices) to *Dmrta2* and *Wnt8b* were reduced from 62 in CTL to 17 in TAM treated cells, and from 109 to 57, respectively, indicating weakened gene-gene interactions in TAM treated cells. These hub genes are critical components of multiple pathways that regulating neural progenitor maintenance, cortical neurogenesis, and patterning (i.e. Wnt, BMP, and Notch signaling pathways) (17, 21).

**Figure 4.**
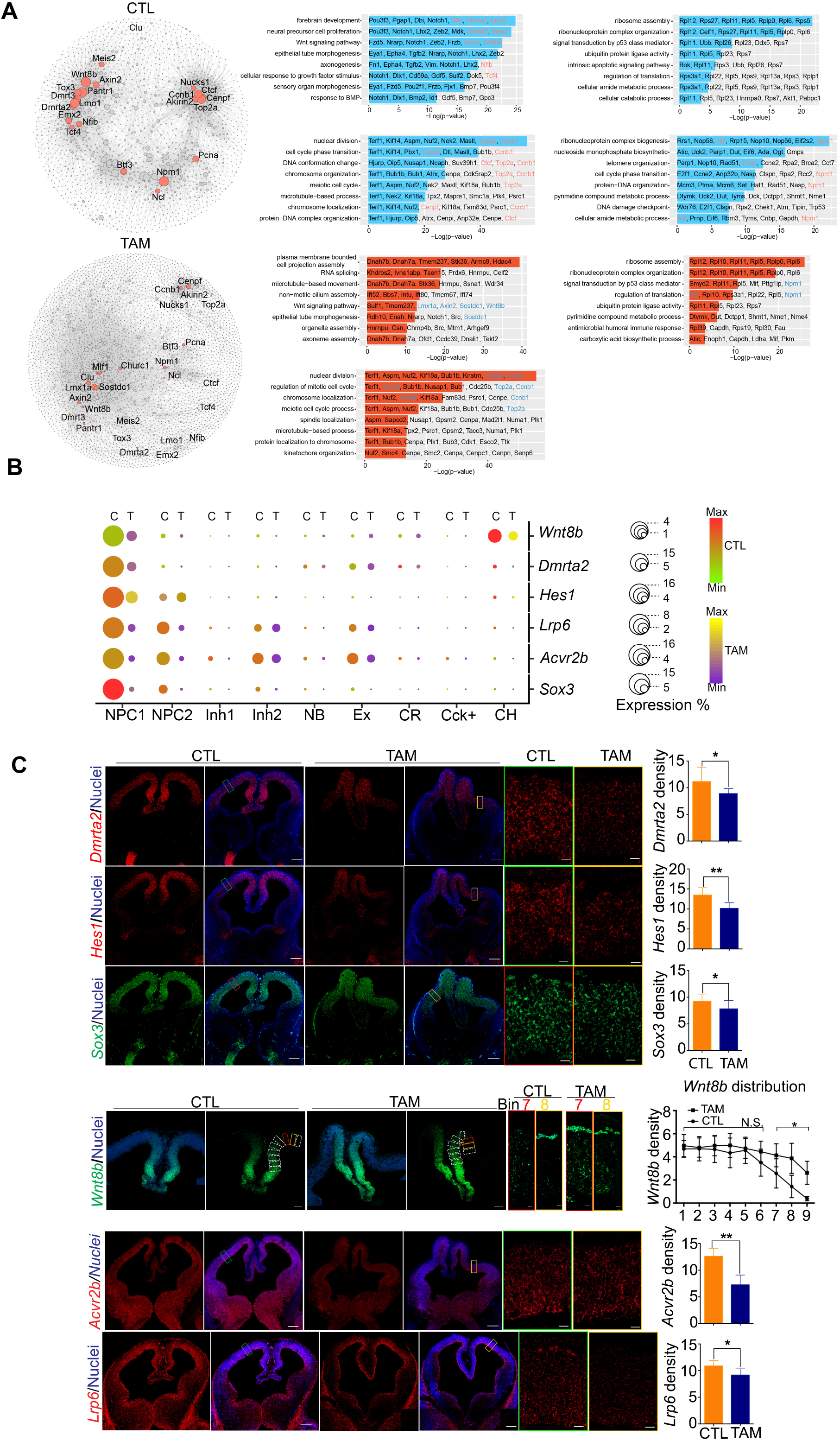
Prenatal TAM administration perturbed Dmrta2-Hes1, Wnt, and Bmp signaling pathways. **(A)** The left panels show CTL and TAM gene regulatory network analysis of scRNA-seq of NPC and CH clusters. The node size is proportional to its pagerank centrality. The orange-red dots show genes in the intersection of three gene sets: 1) top 200 differentially expressed genes identified using Seurat package *FindAllMarkers* function between CTL and TAM; 2) top 200 genes with the biggest change in pagerank absolute value between CTL and TAM; 3) genes with over 0.95 quantile of pagerank centrality. The right panels are GO enrichment of genes with the most affected pagerank centrality. **(B)** The Dotplot shows prenatal TAM administration reduced the number of cells expressing *Dmrta2*, *Hes1*, *Sox3, Lrp6*, or *Acvr2b*. **(C)** ISH assays show reduced number of cells expressing *Dmrta2*, *Hes1*, and *Sox3* in TAM treated brains (*Dmrta2*, p=0.0329, CTL n=12, TAM n=12; *Hes1*, p<0.0001, CTL n=15, TAM n=15; *Sox3*, p=0.0318, CTL n=12, TAM n=12). Scale bars, 200μm and 20μm (zoomed in images). TAM treatment extended *Wnt8b* expression laterally in the cortex (p=0.0009~<0.0001, 2-way ANOVA, CTL n=9, TAM n=9). The cortex above the CH was separated by continuous bins and ISH staining puncta were quantified in each bin. The zoomed in images on the right showed bin-7 and −8. N.S., non-significant. Scale bar, 100μm and 10μm (zoomed in images). TAM treatment decreased numbers of cells expressing a Wnt receptor subunit, *Lrp6* (p=0.0117, CTL n=8, TAM n=8) and a Bmp receptor subunit, *Acvr2b* (p=0.0008, CTL n=13, TAM n=8). Scale bar, 200μm and 20μm (zoomed in images). Error bars, standard deviation.

Intriguingly, a recent study by Young et al. showed that *Dmrta2* regulated *Hes1* and other proneural genes to maintain NPC fate. Emx1-cre conditional knockout of *Dmrta2* accelerated cell cycle exit, increased Eomes+ cells, and promoted transient neuronal differentiation (17). In line with this evidence, an earlier study by Nakamura et. al. showed that *Hes1* null mice exhibited premature progenitor cell differentiation and reduction of multipotent progenitors’ self-renewal activity (16). These phenotypes somewhat resemble our findings in TAM treated brains. scRNA-seq analysis showed that the expression levels of *Dmrta2*, *Wnt8b*, a *Dmrta2* downstream target *Hes1*, a Wnt receptor subunit *Lrp6*, and a Bmp receptor subunit, *Avcr2b* were decreased in the TAM treated NPCs and/or CH cells (Fig. 4B). We further performed ISH assays to show that TAM treatment at E10 dramatically downregulated expression of *Dmrta2* and *Hes1* in the cortex, the cortical hem, and the hippocampus anlage at E12 (Fig. 4B, C). We also verified the reduced expression of a neural progenitor marker, *Sox3* in TAM treated brains (Fig. 4B, C) (30). These results strongly suggested that prenatal TAM administration could trigger precocious cortical neural differentiation and suppress neural progenitor maintenance via suppressing the expression of *Dmrta2* and *Hes1*.

Along with neural progenitor fate maintenance, *Dmrta2*, a novel Wnt-dependent transcription factor, has been shown to be involved in cortical patterning (31, 32). By using a conventional *Dmrta2* null mouse line, Saulnier et at. showed that in the major telencephalic patterning centers, the roof plate and cortical hem, loss of *Dmrta2* resulted in decreased expression of Wnt and BMP signaling genes, such as *Wnt8b* (31). Our scRNA-seq analysis showed TAM-treatment at E10 dysregulated the expression of Wnts, Bmps, receptors, and cytoplasmic modulators in NPCs and cells in the cortical hem (Fig. 4A-C, Sup. Fig. 4A). We found TAM treatment caused expansion of *Wnt8b* expression from ventral-medial to dorsal-lateral region (Fig.4C). TAM also downregulated the expression level of a Wnt receptor subunit, *Lrp6* and a Bmp receptor subunit, *Avcr2b* in the cortex (Fig. 4C). In addition to above genes, the single cell transcriptomic analysis showed TAM also perturbed the expression of genes that have been shown to be involved in Wnt, BMP, Fgf, Notch, and Shh pathways, such as frizzled class receptor 2 (*Fzd2*), glycogen synthase kinase 3 Beta (*Gsk3b*), Cyclin D1, D2, D3 (*Ccnd1-3*), inhibitor of DNA binding 4, HLH protein (*Id4*), protein tyrosine phosphatase non-receptor type 11 (*Ptpn11*), Hes family BHLH transcription factor 5 (*Hes5*), SNW domain containing 1 (*Snw1*), c-terminal binding protein 1 (*Ctbp1*), histone deacetylase 1 (*Hdac1*), *Hdac2*, Gli family zinc finger 3 (*Gli3*), and protein kinase cAMP-dependent type I regulatory subunit alpha (*Prkar1a*) (Sup. Fig. 4A). This evidence suggests that in addition to Wnt8b-Dmrta2-Hes1 axis of regulation, TAM could also exert its effects on NPC fate maintenance and specification through other signaling molecules.

In summary, our data revealed that TAM administration altered the landscape of gene regulatory network in NPCs and CH cells by reducing intra-modular connectivity and gene expression heterogeneity. Particularly, TAM administration at the early stage of cortical neurogenesis dramatically attenuated Wnt signaling network. As a proof-of-principle, we showed a tight link between TAM and Wnt8b-Dmrta2 regulatory axis in regulating NPC fate specification. TAM treatment dysregulates this pathway and leads to deficits in cortical neurogenesis and patterning in prenatal and postnatal offspring.

### TAM regulated cell proliferation and differentiation

To test whether or not TAM can directly regulate cell proliferation and differentiation, we performed pair-cell assay *in vitro* (33). The E11 cortical progenitors were isolated and treated with a TAM active metabolite, 4-hydroxy-tamoxifen (4-OH-TAM). After 16hr, neurons and neural progenitors were detected by immunostaining. The pair-cell assay showed that 4-OH-TAM produced more neurons (Tubb3 (Tuj1)+ cells, N/N) than the control but reduced the numbers of progenitor pairs (P/P) and total Mki67+ cells (Sup. Fig. 5). The *in vitro* assay showed TAM can directly regulate progenitor cell proliferation and differentiation.

**Figure 5.**
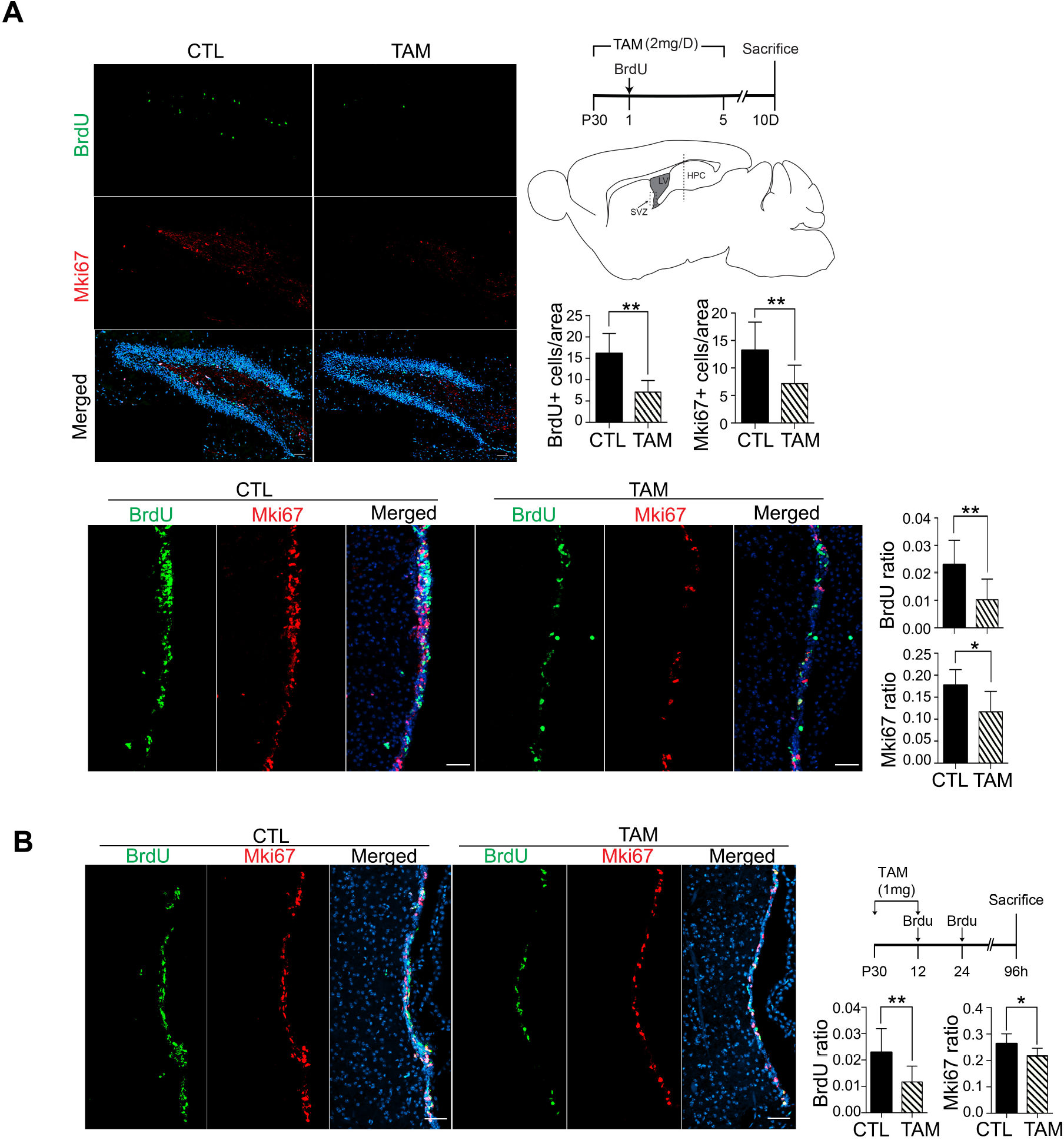
TAM inhibits adult neural stem cell proliferation in both the SVZ of the forebrain and the DG of the hippocampus. **(A)** Mice were given 2mg TAM per day for 5 consecutive days at P30. BrdU was administrated on the second day of TAM injection. Samples were collected 10 days after the initial TAM administration (upper right panel). TAM treatment dramatically decreased number of cells expressing either BrdU or Mki67 in the SVZ (BrdU, p=0.0187, Mann-Whitney test, CTL n=9; TAM n=6; Mki67, p=0.0021, Mann-Whitney test, CTL n=12; TAM n=12) and in the DG (BrdU, P<0.0001, Mann-Whitney test, CTL n=12 TAM n=12; Mki67, p=0.0050, Mann-Whitney test, CTL n=12; TAM n=12). Scale bar, 50μm. LV, lateral ventricle. Scale bar, 100μm. Error bars, standard deviation. **(B)** Mice were given two TAM injections (1mg/mouse) on the first day at P30. The BrdU was administrated 12hr and 24hr after the first TAM injection. Samples were collected 96hr after the initial TAM administration. TAM treatment decreased the numbers of BrdU+ and Mki67+ cells in the SVZ (BrdU, p<0.0001, Mann-Whitney test, CTL n=36; TAM n=31; Mki67, p=0.0401, Mann-Whitney test, CTL n=36; TAM n=31). Scale bar, 50μm.

### TAM inhibited adult neural progenitor cell proliferation in both the SVZ of the caudate putamen and the DG of the hippocampus

In adult mice, several studies reported acute but not long-lasting effects of TAM treatment on locomotion, exploration, and anxiety behaviors, and learning and memory in mice (34-37). Evidently, neurogenesis is disrupted with prenatal administration of TAM. Therefore, would postnatal administration of tamoxifen negatively impact adult neural stem cell proliferation? To study the potential influence of TAM in adult neurogenesis, we employed an established 5 days protocol that was used for CreER/LoxP dependent gene targeting in adult mice (38). We injected TAM (2mg/day) to male mice (C57BL/6) at P30 for continuous 5 days. BrdU was administrated on the second day of TAM injection. The brains were analyzed 5 days after the last TAM administration. We showed that TAM greatly reduced the number of BrdU and Mki67 labeled proliferating cells in the SVZ and in the DG (Fig. 5A). To test the TAM dosage effect, we reduced the TAM dose to 1mg/animal per day and the frequency of TAM treatment to two injections. The mice received two BrdU injections 12hr and 24hr after the first TAM administration. The brain samples were analyzed 96hr after the initial TAM administration. We observed the same effect of reduced proliferating cells in the SVZ of the caudate putamen as in high TAM dosage group (2mg/day) (Fig. 5B). There was no detectable cell apoptosis after 5 days of TAM treatment (2mg/animal/day) (Sup. Fig. 6A) and administration of TAM did not significantly change overall dendritic complexity in adult brains (Sup. Fig. 6B).

## Discussions

Estrogens have been shown to promote NPC proliferation and differentiation, neurite elongation, and synaptic formation during the later stage of corticogenesis (39-41). The current study revealed prenatal, low-dosage administration of an estrogen receptor modulator, TAM, promoted transient SVZ progenitor proliferation, led to precocious cortical neurogenesis, while inhibited VZ progenitor proliferation which in turn severely impaired cortical neurogenesis, cortical patterning, and dendritic morphogenesis in neonatal and postnatal offspring. It seems that TAM and estrogen treatments showed somewhat opposite phenomena in terms of NPC proliferation and differentiation, suggesting TAM may function through interfering with estrogen signaling pathway. The single cell transcriptomic analysis showed that expression of major estrogen receptors, estrogen receptor 1 (*Esr1*), *Esr2*, G Protein-Coupled Estrogen Receptor 1 (*Gper1*), estrogen-related receptor beta (*Esrrb*), and a critical estrogen synthetic enzyme, aromatase (*Cyp19a1*) were either extremely low or undetectable at E12 in the telencephalon, while the expression of estrogen-related receptor alpha (*Esrra*) was low and TAM treatment at E10 did not alter its expression at E12 (Sup. Fig. 4B). Along with the evidence that 4-OH-TAM could directly regulate NPC proliferation and differentiation (Sup. Fig. 5), it seems that at the early stage of cortical neurogenesis (E10-12) TAM may directly interfere with normal cortical development via Esrra and/or Gper1 or estrogen receptor-independent mechanisms (42). In the meantime, the fact that a single dosage of prenatal TAM administration at early stage of brain development causes dystocia, indicates that the effect of TAM on pregnant dams can be long lasting and TAM exposure may indirectly influence cortical neurogenesis of fetuses.

The single cell transcriptomic analysis substantiates that prenatal TAM treatment altered gene-to-gene regulatory networks which regulate forebrain development, chromosome replication, cell cycle progression, and translation regulation in NPC and the cortical hem clusters. We showed that TAM administration drastically inhibited NPC proliferation, and thus led to impairment of cortical neurogenesis and patterning in neonatal and postnatal offspring. Mechanistically, we showed that prenatal TAM administration downregulated the expression of a Wnt related transcription factor *Dmrta2* and its effector *Hes1*. Dmrta2 has been shown to play critical roles in NPC fate maintenance and differentiation (16, 17). TAM also downregulated the expression of Wnt and BMP receptors, *Lrp6* and *Avcr2b* in cortical NPCs. TAM induced perturbation of Wnt and BMP pathways could somewhat explain the underlying mechanism of cortical patterning changes in TAM treatment neonatal brains. Additionally, downregulation of *Emx2*, *Dmrt3*, and *Gli3* expression may also contribute to cortical neurogenesis and patterning deficits in TAM treated brains (Fig. 4A, Sup. Fig. 4A) (31). Moreover, the single cell transcriptomic analysis showed dramatical reductions of Wnt signaling-related genes (e.g. *Fzd2*, *Gsk3b*, *Ccnd1,2,3*), Tgf-β pathway gene (*Id4*), Fgf pathway gene (*Ptpn11*), Notch pathway genes (*Hes5*, *Snw1*, *Ctbp1*, *Ctbp2*, *Hdac1*, *Hdac2*) and sonic hedgehog (Shh) pathway genes (*Prkar1a*, *Smo*) in NPCs (Sup. Fig. 4A). Taken together, these data suggest that in addition to the Wnt8b-Dmrta2-Hes1 axis of regulation, other signaling molecules that could also be involved in TAM regulation of NPC fate maintenance and specification during corticogenesis, as well as dendritic morphogenesis and axon formation.

TAM inducible CreER/LoxP system laid the foundations of significant discoveries in adult and embryonic neural stem cell fate mapping (2, 10, 11, 43), gene functions (44), as well as neuronal subtypes and activity dependent neural circuitry (45-47). Is the potential side effect of TAM a vulnerable Achille’s heel to TAM-induced CreER/loxP cell lineage tracing and/or genetic targeting studies? Now, it is clear that prenatal, single dosage of TAM exposure dramatically changes the global gene expression landscapes of cells in the cerebral hemisphere, which has long lasting influence on cortical neurogenesis and patterning in perinatal and postnatal offspring. Therefore, special cautions should be taken when using TAM-induced CreER/LoxP system for neural lineage tracing study because it is not falsifiable due to lack of appropriate controls.

## Materials and Methods

### Experimental Animals

Three different genetic background lines were used: C57BL/6, Crl:CD1(ICR), and Prom1creER/ZsGreen transgenic line, which was generated by crossing Prom1tm1(cre/ERT2)Gilb line (129S6/SvEvTac background) and Gt(ROSA)26Sortm6(CAG-ZsGreen1)Hze (129S6/SvEvTac x C57BL/6NCrl background, JAX) (Details are given in SI Materials and Methods).

### TAM administration and fostering

TAM (Sigma, T5648) was suspended in corn oil to a stock concentration of 20mg/ml. Tamoxifen was administrated by IP injection. To obtain postnatal offspring, a caesarean section was performed on pregnant mothers in the afternoon on E18.5 and pups were fostered to CD1 or 129S6 mothers (Details are given in SI Materials and Methods).

### Single cell transcriptomic analysis

The forebrains were collected at E12, dissociated in papain (BrainBits, LLC). The cell viability was accessed by trypan blue staining. The cells were diluted to a final concentration of 1×10^6^/ml in PBS with 0.04% BSA. The volume of single cell suspension that was required to generate 10,000 single cell GEMs (gel beads in emulsion) per sample was loaded onto the Chromium Controller (10x Genomics).

The raw reads were processed to molecule counts using the Cell Ranger pipeline (version 2.1.1, 10X Genomics). The raw unique molecular identified (UMI) counts from Cell Ranger were processed with the Seurat R toolkit (version 3.0.1). The cell numbers of CTL and TAM datasets were then normalized by random section of 4,193 cells/condition. The top 2,000 genes that exhibited high cell-to-cell variation in each dataset (CTL, TAM) were identified respectively via *FindVariableFeatures* function. Based on UMAP plot, fourteen clusters were classified using the function *FindClusters* with resolution parameter set to 0.4. To identify the marker genes of each cluster, function *FindAllMarkers* with likelihood-ratio test was applied. According to expression of well-established marker gene sets, the 14 clusters were manually annotated as major classes of cells.

The NPC1, NPC2, and CH clusters of each condition (CTL, TAM) were merged and served as the input for inferring the putative gene-to-gene correlated network using bigSCale2 algorithm (28). Specifically, two matrices with 17,071 genes’ expression counts for 1,908 CTL and 1,127 TAM cells were used to infer the network by the default parameters (i.e. clustering = ‘recursive’, quantile.p = 0.998). The network centrality pagerank was chosen to represent the gene essentiality (Details are given in SI Materials and Methods).

### Studying cell cycle with 5-ethynyl-2’-deoxyuridine (EdU) and bromodeoxyuridine (BrdU)

The pregnant mice were treated with TAM at E11. At E12, EdU (100mg/kg, Abcam ab219801) was IP delivered to the TAM treated mice followed by BrdU (100mg/kg) labeling after 2hr. Details are given in SI Materials and Methods.

### BrdU administration

Details are given in SI Materials and Methods.

### *In vitro* TAM assay

Telencephalons of E11 CD-1 embryos were collected and dissociated single cells treated with 1µM 4-OH-TAM (Sigma, H6278). 16hr after plating, the cells were fixed with 4% PFA at room temperature. Details are given in SI Materials and Methods.

### In situ hybridization (ISH)

Fluorescent ISH assay was performed followed manufacturer’s instructions (Advanced Cell Diagnostics, Inc.). Details are given in SI Materials and Methods.

### Immunohistochemistry

The primary antibodies used in this study were BrdU, MKi67, phospho-Histone H3 (Ser10), Map2, Tbr1, Pax6, Emoes, Rbfox3, Bcl11b, Mef2c, Mbp, Cnp, Olig2, Aldoc, and Gfap. Details are given in SI Materials and Methods.

### Golgi-Cox Staining

The Golgi staining was carried out following manufacturer’s instructions (IHCWORLD, IW-3023).

### Imaging and Statistical Analysis

Images obtained were processed and quantified with Imaris software. Statistical analysis of the data was carried out with Prism 6. Details are given in SI Materials and Methods.

## Acknowledgements

We thank the Intellectual and Developmental Disabilities Research Center (IDDRC) at the University of California, Los Angeles (UCLA) supported by NIH Grant U54HD087101. This work was supported by NIH/National Institute of Mental Health (NIMH) Grant 5R21MH115382 (Q.L. and Y.E.S).

## Author contributions

Q.L., Y.E.S. CM.L; LQ. Z. designed research; CM.L., LQ. Z., JP.L., JY.S., YN.G. JR.W., XJ.S., N.B., JB.W. performed research and scRNA-seq data analysis; Q.L, CM.L., LQ.Z. and Y.E.S. wrote the paper.

## Notes

#### Summary of Updates

corrected some errors in the manuscript.

