## Supplementary material for "Single cell transcriptomic analysis revealed long-lasting adverse effects of prenatal tamoxifen administration on neurogenesis in prenatal and adult brains": Materials and Methods

**Supplementary information**

**Materials and Methods**

**Experimental Animals.** Mice were kept and fed in standard conditions on a 12hr light/dark cycle. Experimental procedures on animals were performed in accordance with the guidelines of UCLA Institutional Animal Care and Use Committee and UCLA Animal Research Committee. Three different genetic background lines were used: C57BL6/J, Crl:CD1(ICR), and Prom1creER/ZsGreen transgenic line, which was generated by crossing Prom1$_{tm1(cre/ERT2)Gilb}$ line (129S6/SvEvTac background, the Jackson Laboratory (JAX)) and Gt(ROSA)26Sor$_{tm6(CAG\text{-}ZsGreen1)Hze}$ (129S6/SvEvTac x C57BL/6NCrl background, JAX). Pregnancy was timed by daily monitoring for vaginal plugs.

Prenatal and perinatal brains were dissected without perfusion, and subsequently immersed overnight in 4% paraformaldehyde (PFA) in PBS. Adult mice were overdosed with isoflurane and perfused intracardially with PBS followed by 4% PFA. The brains were removed and stored in 4% PFA overnight at 4 °C. Fixed prenatal and perinatal brains were coronally sectioned at 10-12μm for BrdU immunostaining and ISH, or at 20-60μm for free-floating immunostaining. Fixed adult brains were sagittal sectioned at 10μm for BrdU/Mki67 immunostaining and TUNEL assay.

**TAM administration and fostering.** TAM (Sigma, T5648) was suspended in corn oil to a stock concentration of 20mg/ml. The injection site was cleaned with ethanol. 750μg of tamoxifen was injected into the peritoneal cavity (Intraperitoneal injection, IP) with 1ml, 28G1/2 insulin syringe. Animals were monitored until proper post injection recovery was demonstrated. Prenatal administration of TAM caused mortality of pups was always observed in more than 60 TAM treated pregnant dams, which were injected with TAM at E8, E10, and E13. To circumvent this problem, a caesarean section was performed on pregnant mothers in the afternoon on E18.5 and pups were fostered to CD1 or 129S6 mothers.

TAM was injected to C57BL/6J mice for 2 (1mg/mouse) or 5 days (2mg/mouse) at P30.

**Single cell transcriptomic analysis.** 750μg of TAM was injected to pregnant mice at embryonic day 10 (E10). The forebrains were collected at E12, dissociated in papain at a concentration of 2 mg/ml in Hibernate E-Ca (HE-Ca) (BrainBits, LLC.) at 30°C for 10min. The digested tissues were gently triturated 10 times with fire-polished pasture pipettes. The single cell suspension was filtered through a 40μm cell strainer, and then centrifuged at 200g for 5min at room temperature. The pellet was washed with PBS with 1% bovine serum albumin (BSA). The cell viability was accessed by trypan blue staining. The cells were diluted to a final concentration of $1x10_6$/ml in PBS with 0.04% BSA. The volume of single cell suspension that was required to generate 10,000 single cell GEMs (gel beads in emulsion) per sample was loaded onto the Chromium Controller

(10x Genomics). Libraries were prepared using the Chromium v2 Single Cell 3′ Library and Gel Bead Kit (10x Genomics) according to the manufacturer's specifications. Final library quantification and quality check was performed using a DNA 1000 chip (Agilent Technologies), followed by sequencing on Illumina NovaSeq 6000.

The raw reads were processed to molecule counts using the Cell Ranger pipeline (version 2.1.1, 10X Genomics) with default settings. The raw unique molecular identified (UMI) counts from Cell Ranger were processed with the Seurat R toolkit (version 3.0.1). Cells derived from the diencephalon were removed. Genes that were detected in less than 0.1% cells were discarded. Low-quality cells that had over 5% mitochondrial UMI counts were removed. Additionally, cells that contained under 400 gene counts or above 8100 gene counts, in the CTL dataset or cells contained either less than 500 gene counts or more than 7,000 gene counts in the TAM dataset were removed. The cell numbers of CTL and TAM datasets were then normalized by random section of 4,193 cells/condition. The gene expression for each cell was then normalized with default parameters. The top 2,000 genes that exhibited high cell-to-cell variation in each dataset (CTL, TAM) were identified respectively via *FindVariableFeatures* function.

A list of CTL and TAM Seurat objects served as the input to identify anchors, which were subsequently used to integrate the two datasets using the function *IntegrateData*, with dims set to 18. Then the expression of each gene was scaled and unwanted sources of variation (UMI counts, mitochondrial contamination) were regressed out via *ScaleData* function. The first 20 principal components (PCs) were chosen to perform principal component analysis (PCA) through function *RunPCA*, followed by shared nearest neighbor (SNN) graph construction via *FindNeighbors* function and uniform manifold approximation and projection (UMAP) dimension reduction using function *RunUMAP*. Based on UMAP plot, fourteen clusters were classified using the function *FindClusters* with resolution parameter set to 0.4. To identify the marker genes of each cluster, function *FindAllMarkers* with likelihood-ratio test was applied. According to expression of well-established marker gene sets, the 14 clusters were manually annotated as major classes of cells: neuronal cells (including pallial neural progenitors, subpallial neural progenitors, neuroblasts, excitatory neurons, inhibitory neurons, Cajal-Retzius cells, Cck+ neurons, and cortical hem cells), and non-neuronal cells (consisting of neural crest cells, red blood cells, endothelial cells, microglia, and pericytes).

The fractions of major types of cells in each dataset were calculated as the percentage of number of each cell type in total cell number of each dataset. The positive ratio of a specific gene in neuronal cells was defined as the ratio of gene positive cells (with log2 transferred UMI counts greater than 0) in each dataset.

The NPC1 (pallial neural progenitors), NPC2 (subpallial neural progenitors), and CH (cortical hem) clusters of each condition (CTL, TAM) were merged and served as the input for inferring the putative gene-to-gene correlated network using bigSCale2 algorithm (1). Specifically, two matrices with 17,071 genes' expression counts for 1,908 CTL and 1,127 TAM cells were used to infer the network by the default parameters (i.e. clustering = 'recursive', quantile.p = 0.998). The network centrality pagerank was chosen to represent the gene essentiality. The network was then visualized via igraph R package (version 0.7.1), with layout setting layout.fruchterman.reingold. Communities in each network were discovered using *cluster_label_prop* function (igraph 1.2.4.1), and clusters consisting of over 100 genes were determined as main communities. The genes in each main community were set as input for gene ontology (GO) enrichment analysis using the R package clusterProfiler (version 3.12.0).

**Bromodeoxyuridine (BrdU) administration.** BrdU (Sigma, B5002) was suspended in water to a working concentration of 10mg/ml. Prior to injection, animals were weighed. After cleaning injection site with ethanol, BrdU solution was administered through IP injection at 100mg/kg (pregnant dams) and 200mg/kg (adult mice). Animals were monitored until proper post injection recovery was demonstrated.

**Studying cell cycle with 5-ethynyl-2'-deoxyuridine (EdU) and BrdU.** The pregnant mice were treated with TAM at E11. At E12, EdU (100mg/kg, Abcam ab219801) was IP delivered to the TAM treated mice followed by BrdU (100mg/kg) labeling after 2hr. The embryonic brains were fixed with 4% PFA after 30min. The cells in the initial EdU-labeled cohort that left S-phase during the interval (Ti) between EdU and BrdU (2hr) were labeled with EdU but not BrdU (the leaving fraction, L). The proportion of cells labeled with BrdU is designated S. The length of S-phase (Ts) can be calculated using the formula: Ts/Ti=S/L.

***In vitro* TAM assay**. Telencephalons of E11 CD-1 embryos were dissected and digested with papain in HBSS at 30°C for 5-10min. The digested tissue was dissociated using fire-polished pasture pipettes. Dissociated single cells were collected by centrifugation and resuspended in neurobasal medium (ThermoFisher Scientific, 21103049) supplied with B27 (ThermoFisher Scientific, 17504044), bFGF (Sigma, F3685), EGF (Sigma E4127), and 1μM 4-OH-TAM (Sigma, H6278). The single cell suspension was plated on Poly-L-Ornithine hydrochloride (Sigma, P-2533) and Fibronectin Human Protein (ThermoFisher Scientific, 33016015) coated coverslips in 24-well plates. For controls, dissociated cells were cultured in neurobasal medium supplied with B27, bFGF, and EGF. 16hr after plating, the cells were fixed with 4% PFA at room temperature.

**In situ hybridization (ISH).** Fluorescent ISH assay was performed followed manufacturer's instructions (Multiplex Fluorescent Reagent Kit v2, Advanced Cell Diagnostics, Inc.). RNAscope Probes were *Fezf2* (313301-C3), *Pax6* (412821), *Eomes* (429641-C3), *Rbfox3* (313311-C2), *Cdkn1c* (458331-C2), *Wnt8b* (405071), *Wnt3a* (405041-C2), *Hes1* (417701-C4), *Sox*3 (customized probe), *Dmrta2* (customized probe), *Acvr2b* (469791-C3), *Lrp6* (315801-C2), *Mdga1* (546411), *Cyp26b1* (524001-C3), *Crym* (466131-C3), and *Epha7* (430961-C2).

**Immunohistochemistry.** The primary antibodies used in this study were BrdU (Abcam ab6326, 1:100; ab220076 1:300), MKi67(Abcam ab15580, 1:350), phospho-Histone H3 (Ser10) (Cell Signaling 9701S,

1:300), Map2 (Abcam ab92434, 1:350), Tbr1 (Abcam ab31940, 1:1000), Pax6 (LSbio LS-C179903, 1:100), Emoes (Abcam ab183991, 1:500 and EMD Millipore ab15894, 1:250), Rbfox3 (EMD Millipore MAB377, 1:50 and Invitrogen 7011084, 1:200, and Encor CPCA-FOX3, 1:500), Bcl11b (Abcam ab18465 (25B6), 1:500), Mef2c (Genetex GTX-105433, 1:750), Mbp (Myelin basic protein) (Encor CPCA-MBP, 1:5,000), Cnp (2', 3'-cyclic nucleotide 3'-phosphodiesterase) (Encor MCA-1H10, 1:1,000), Olig2 (EMD Millipore, 1:150), Aldoc (Encor MSA-4A9, 1:200) (2), and Gfap (Abcam ab7260, 1:500). For BrdU/Mki67 staining, sections were first stained with Mki67. After secondary antibody incubation, the sections were post-fixed with 4% PFA for 30 minutes. Then, the sections were treated with 2N HCl for 20 minutes at 37°C, neutralized with 0.5% sodium borate buffer, and incubated overnight with BrdU.

Fluorophore-conjugated secondary antibodies were purchased from ThermoFisher Scientific and Jackson ImmunoResearch.

**Golgi-Cox Staining.** TAM was IP injected to adult mice (~P60) for 5 days (2mg/mouse) and the brains were removed 14 days after the last TAM injection. The Golgi staining was carried out following manufacturer's instructions (IHCWORLD, IW-3023).

**Imaging and Statistical Analysis.** Stained sections were imaged with Zeiss LSM800 confocal microscope as described in our previous work (3). Images obtained were processed and quantified with Imaris software. Statistical analysis of the data was carried out with Prism 6. Unpaired, two-tailed T-tests with equal standard deviation were used to assess statistical significance between independent experimental groups. All reported significance levels represent two-tailed values.

**Figure legends**

**Supplementary Figure 1. Prenatal TAM exposure impaired neural progenitor proliferation by interfering with cell cycle progression. (A)** To quantify cells that expressed *Pax6*, *Eomes*, or both, the nuclei were marked with the Imaris create surface tool. The *Pax6* and *Emoes* puncta were filtered based on their localization. Only a surface containing more than three puncta was counted as a positive cell. **(B)** Prenatal TAM administration at the peak period of cortical neurogenesis (E13) dramatically reduced the number of cells in the S-phase (BrdU labeled at E14) and total number of proliferating cells (Mki67+) (BrdU, $p=0.0010$, Mki67, $p<0.0001$, Mann Whitney test, CTL n=13, TAM n=18). Samples were collected 30min after BrdU administration. Scale bar, 60μm. **(C)** *Cdkn1c* ISH assay suggested that prenatal administration of TAM at E13 promoted cells entering G1-phase. Scale bar, 30μm. Error bar, standard deviation. **(D)** TUNEL assay shows that prenatal administration of TAM (2mg/animal) did not cause cell apoptosis. Scale bars, 200μm, 30μm (zoomed in images).

**Supplementary Figure 2. Prenatal TAM exposure impaired cortical neurogenesis. (A)** Administration of TAM at the peak period of cortical neurogenesis (E13) did not change the number of Reln+ Cajal–Retzius cells at E14 ($p= 0.5179$, Mann-Whitney test, CTL n=17, TAM n=21). Scale bar, 30μm. TAM transiently increased number of Tbr1+ early born neurons and superficial layer Mef2c+ cells in prenatal brains (Tbr1, $p=0.0002$, Mann-Whitney test, CTL n=20, TAM n=17; Mef2c, $p<0.0001$, Mann-Whitney test, CTL n=20, TAM n=20). Scale bars, 30μm (Tbr1) and 20μm (Mef2c). **(B)** Administration of TAM at the peak period of cortical neurogenesis (E13) reduced the number of Rbfox3+ cortical neurons in E18 brains. Scale bar, 100μm. **(C)** Prenatal administration of TAM during early brain development reduced the number of cortical neurons in an outbred stain ICR CD1 and Prom1creER/ZsGreen transgenic mouse line (129S6/SvEvTac background). Scale bar, 100μm.

**Supplementary Figure 3. The long-lasting effects of prenatal TAM administration on astrogliogenesis in postnatal offspring. (A)** Early prenatal TAM treatment impaired gliogenesis. Olig2-, Mbp-, and Cnp-positive signals decreased in the corpus collosum of TAM treated brain (Olig2+ cells, $p=0.0425$, Mann-Whitney test, CTL n=14; TAM n=14). Scale bar, 100μm. Olig2, oligodendrocyte transcription factor 2; Mbp, myelin basic protein; Cnp, 2'3' cyclic nucleotide 3' phosphodiesterase. **(B, C)** We did not detect a significant alteration in the number of astrocytes in both the corpus collosum (B) and the cortex (C) of E8.5 TAM treated brain (B, Gfap+ cells, $p=0.7198$, Student's t-test, CTL n=14, TAM n=14; C, Aldoc+ cells, $p=0.0592$, Student's t- test, CTL n=18, TAM n=15; ). Gfap, glial fibrillary acidic protein; Aldoc, aldolase, fructose-bisphosphate C. Scale bar, 80μm. Error bars, standard deviation.

**Supplementary Figure 4. Gene expression that was altered by prenatal TAM administration. (A)** Signaling pathways that may be involved in cell proliferation, differentiation, and brain area patterning. The size of dot shows the percentage of cells expressing detectable target genes in each cluster. **(B)** Violin plot combined with box plot shows the overall expression of estrogen signaling related genes.

**Supplementary Figure 5. TAM directly regulated neural progenitor proliferation and neurogenesis. (A)** *In vitro* cell proliferation assay showed that treatment of E11 cortical NPCs with 4-hydroxytamoxifen (4-OH-TAM) significantly increased new born neurons (upper panels, Tubb3+), while decreased number of progenitor cells (middle panels, Nestin+) (N/N: $p=0.0008$; P/P: $p<0.0001$; N/P: $p=0.5360$, Mann Whitney test, CTL n=34, TAM n=33). Scale bars, 5μm(N/N), 2μm(P/P), 3μm(N/P). N, neuron, P, progenitor. Tubb3, neuron-specific class III β-tubulin. **(B)** Treatment of E11 cortical NPCs with 4-OH-TAM dramatically reduced Mki67+ proliferating cells ($p<0.0001$, Mann Whitney test, CTL n=17, TAM n=19). Scale bar, 20μm. Error bars, standard deviation.

**Supplementary Figure 6. Administration of TAM in adult neither caused cell death nor drastically reduced dendritic complexity. (A)** TUNEL assay shows that administration of TAM in adult mice at 2mg/animal for 5 days did not cause cell death in the SVZ and the DG area. CPu, caudate putamen, LV, lateral ventricle, MZ, marginal zone, DG, dentate gyrus. Scale bars, 40μm (cortex), 100μm (DG), 50μm (CPu). **(B)** Golgi staining shows that administration of TAM in adult mice did not dramatically alter dendritic complexity of cortical neurons (p=0.6898, Student's t- test, CTL n=29, TAM n=65). Scale bar, 100μm.
