## Supplemental Figures for "Single cell transcriptomic analysis revealed long-lasting adverse effects of prenatal tamoxifen administration on neurogenesis in prenatal and adult brains"

Supplementary Figure 1 (Lee & Zhou et al.)

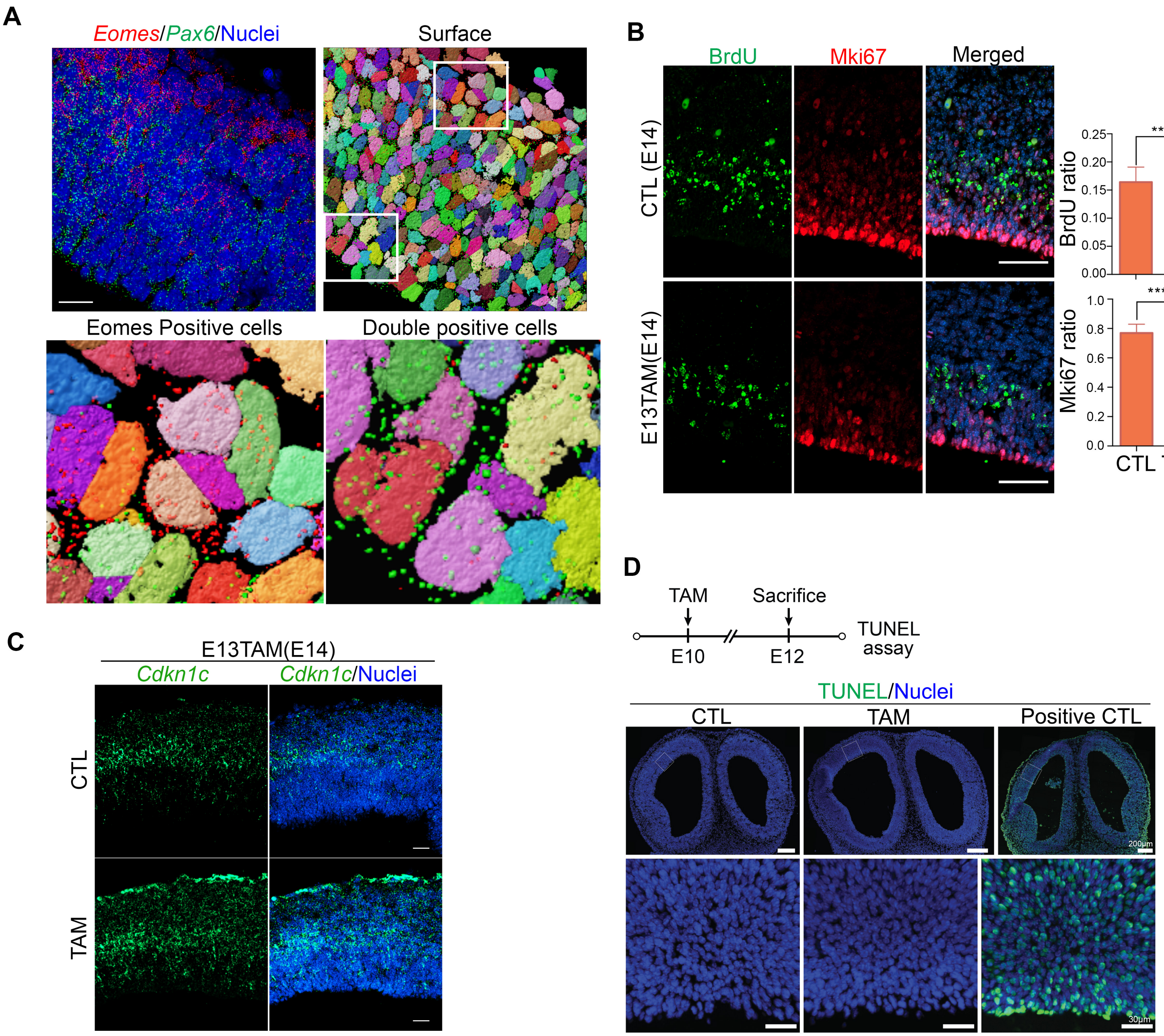

**Supplymentary Figure 2 (Lee & Zhou et al.)**

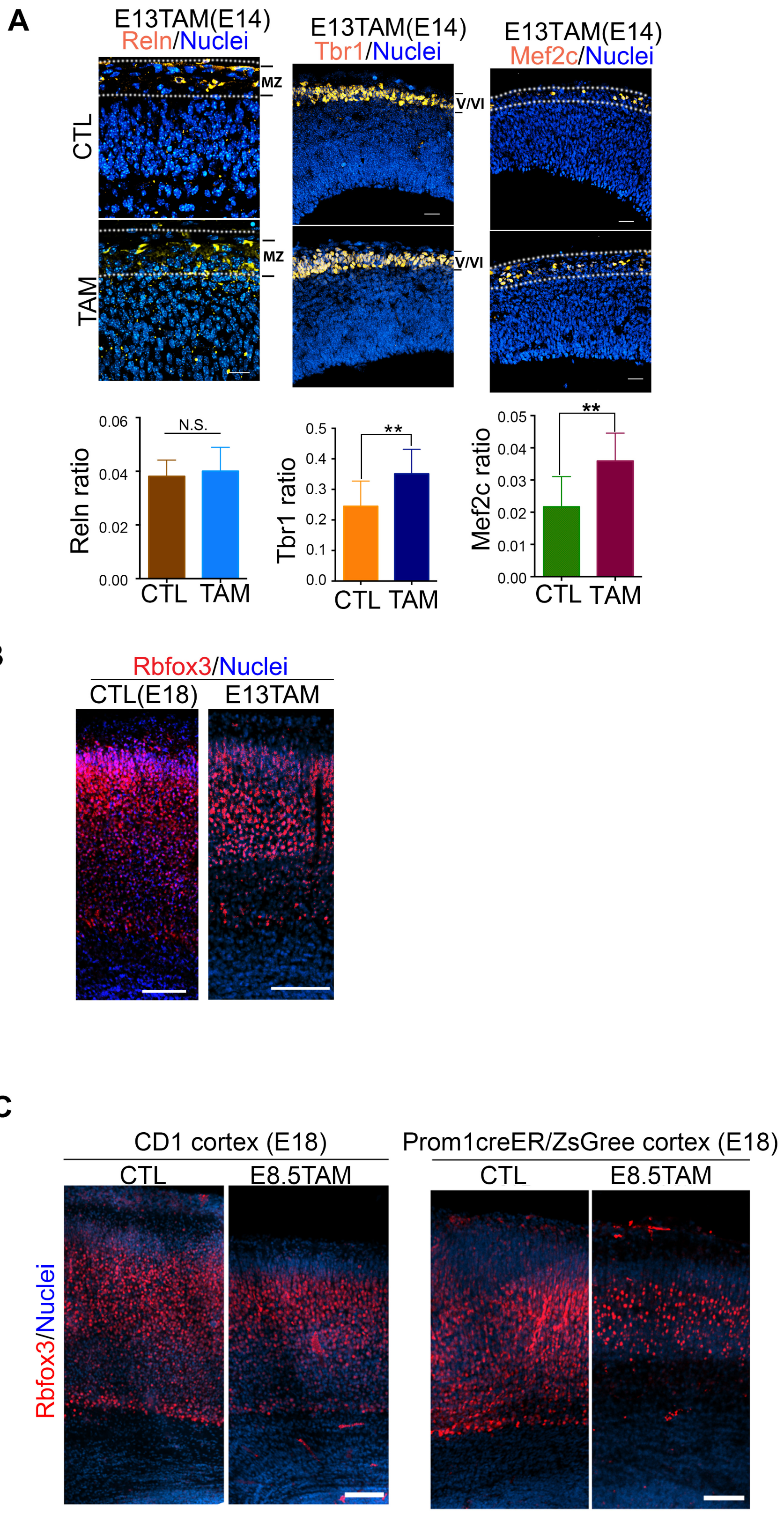

#### Supplementary Figure 3 (Lee & Zhou et al.)

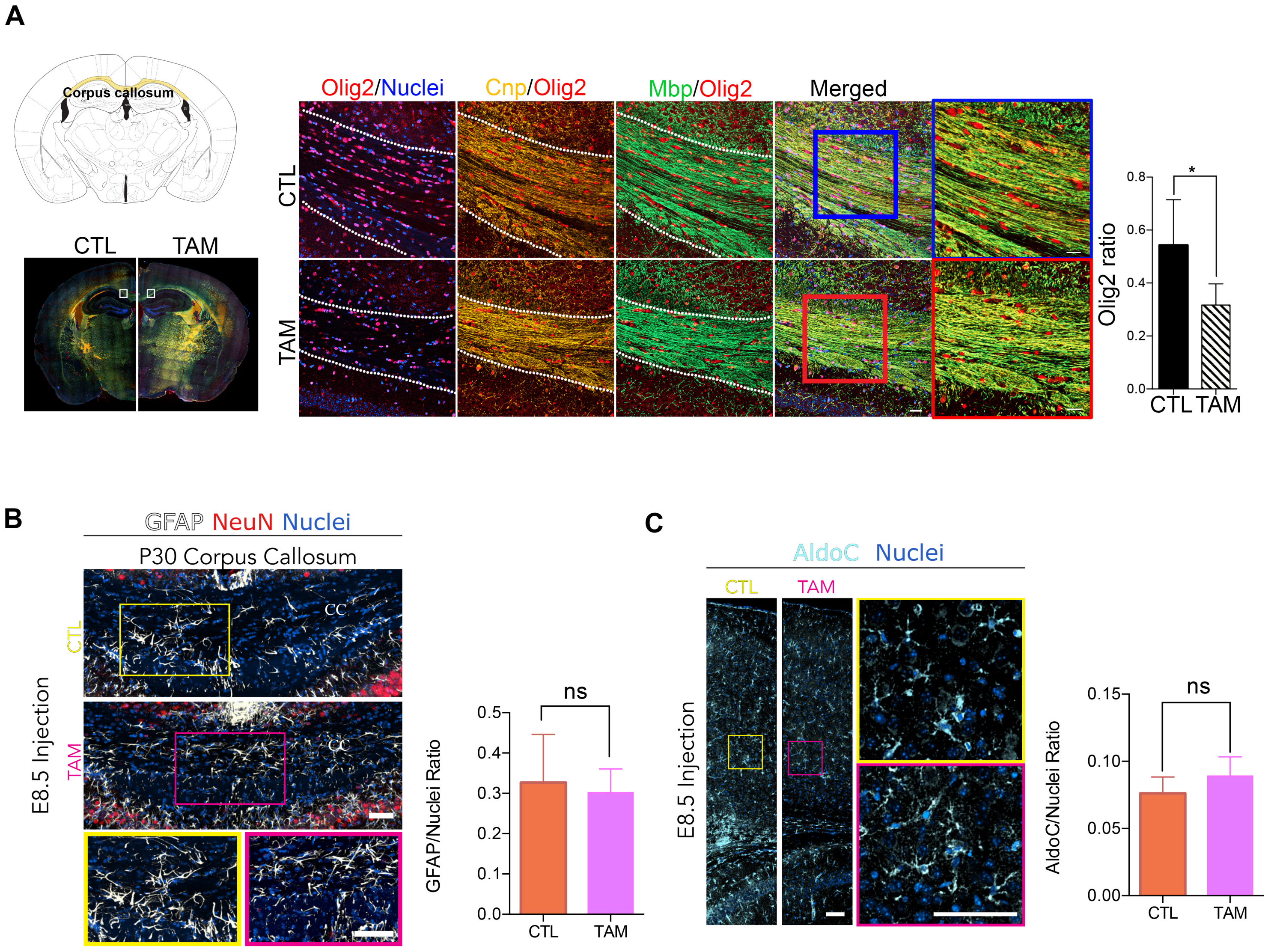

A

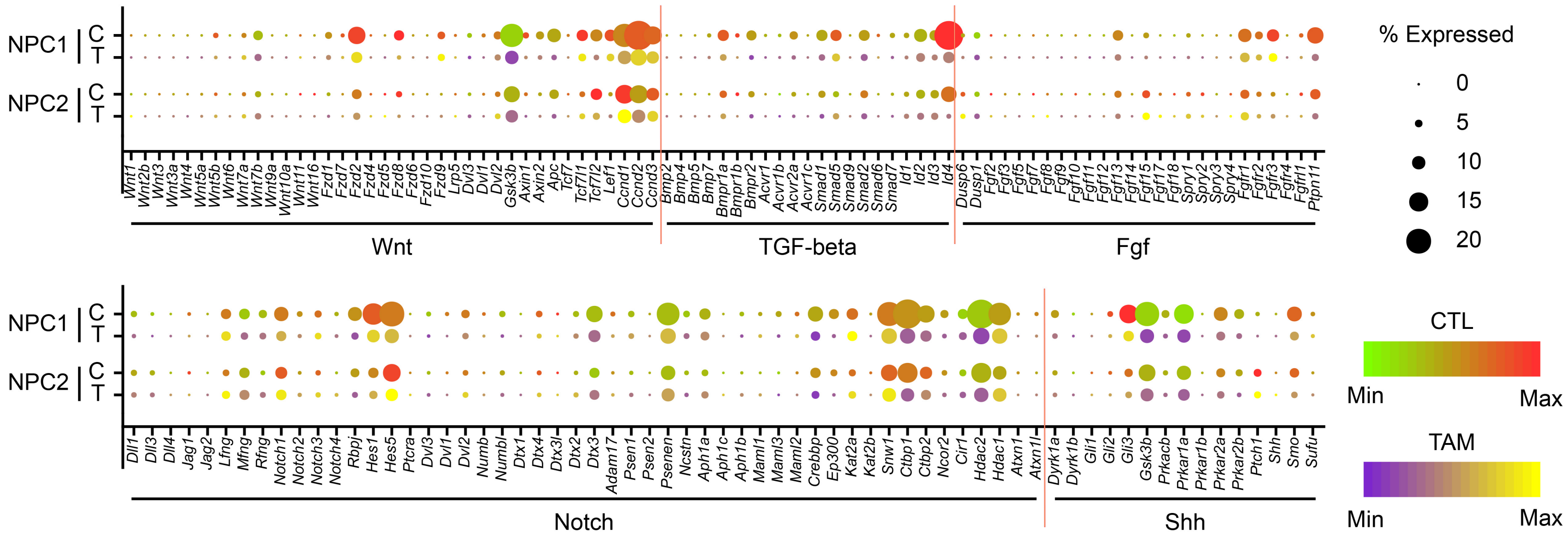
NPC1
C
T
NPC2
C
T
Wnt
TGF-beta
Fgf
Notch
Shh
% Expressed
0
5
10
15
20
CTL
Min
Max
TAM
Min
Max

B

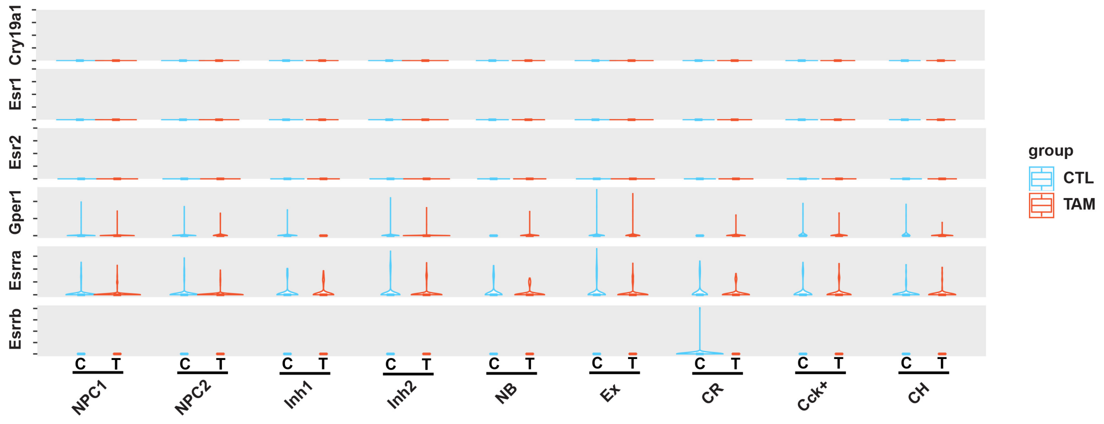
Cry19a1
Esr1
Esr2
Gper1
Esrra
Esrrb
group
CTL
TAM
NPC1
NPC2
Inh1
Inh2
NB
Ex
CR
Cck+
CH

### Supplementary Figure 5 (Lee & Zhou et al.)

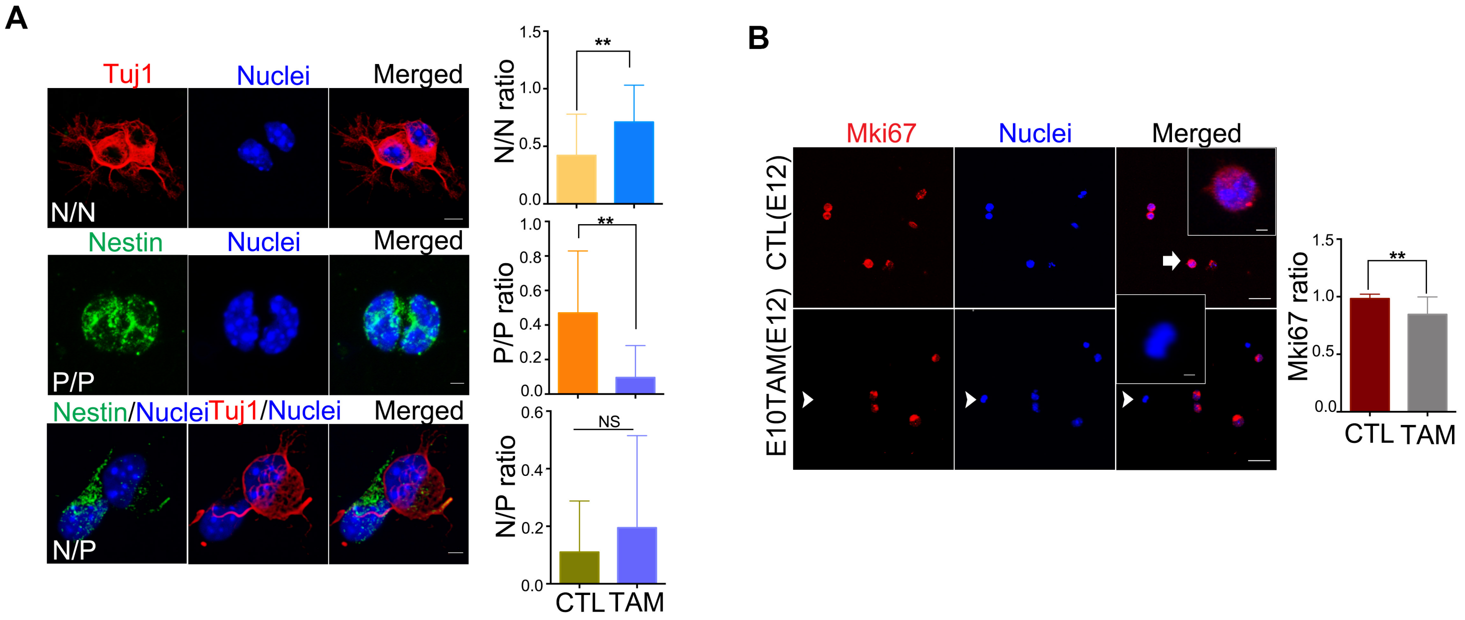

### Supplementary Figure 6 (Lee & Zhou et al.)

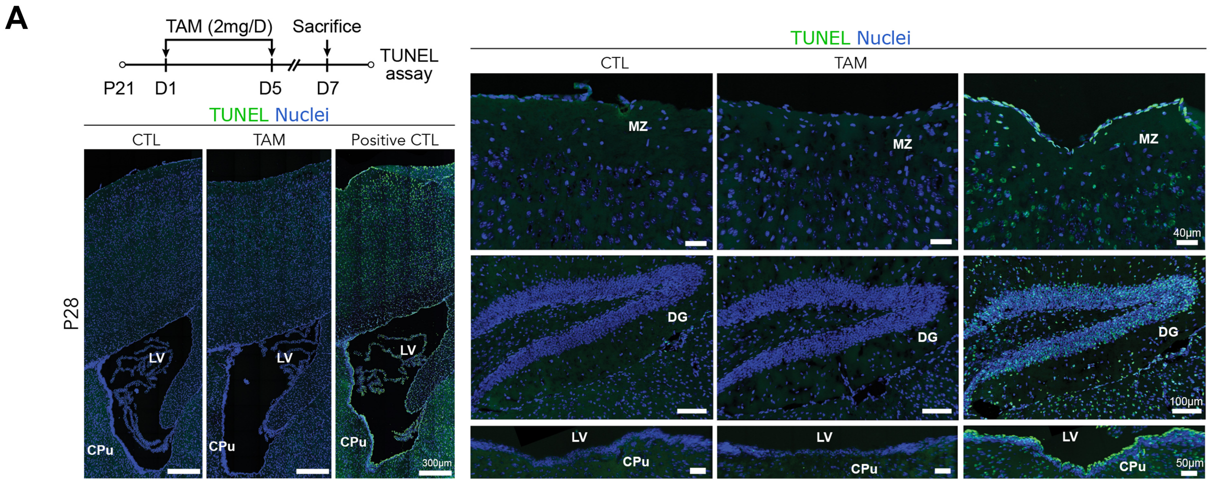


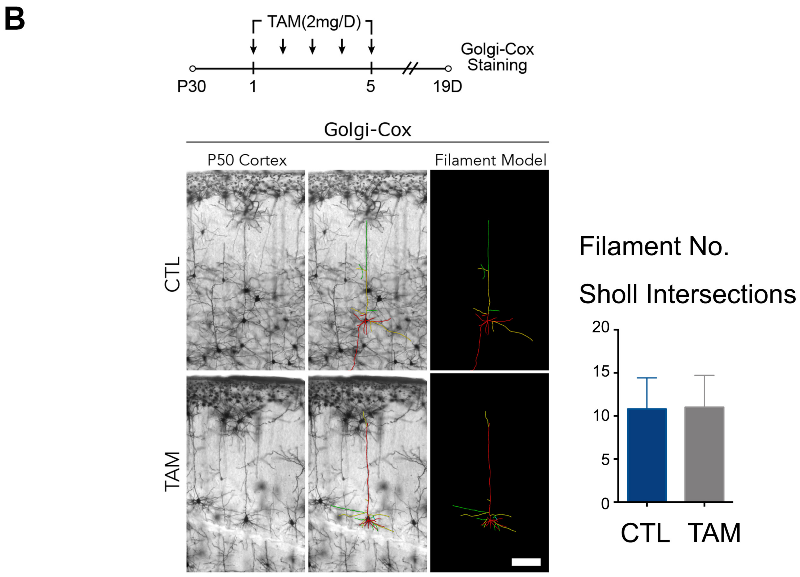
